# Loss of PEA-15 Alters ERK/MAPK Signaling and Circadian Function in the Suprachiasmatic Nucleus

**DOI:** 10.64898/2026.10.25.761398

**Authors:** Kelin Wheaton, Nicklaus Halloy, Megan Formanowicz, Joe W. Ramos, Kari R. Hoyt, Karl Obrietan

**Author notes:** Corresponding Authors: Karl Obrietan Kari R. Hoyt Department of Neuroscience Division of Pharmaceutics and Pharmacology Ohio State University Ohio State University Graves Hall, Rm 4036 Riffe Building, Rm 412 333 W. 10th Ave. Columbus, OH 43210 496 W. 12th Ave. Columbus, OH 43210.

## Abstract

Circadian rhythms are a fundamental biological process that coordinates physiological functions, behavior, and metabolism with an approximately 24-hour periodicity. At the center of the mammalian circadian timing system is the suprachiasmatic nucleus (SCN) of the hypothalamus, which generates robust, clock-gated neuronal output that establishes the temporal organization of cellular and physiological processes throughout the body. Our previous work demonstrated that the scaffold protein PEA-15 (phosphoprotein enriched in astrocytes 15) is highly expressed in the SCN and dynamically interacts with ERK, a key effector kinase of the MAPK pathway, to regulate its subcellular localization. Given the important role of ERK/MAPK signaling in circadian timing and photic entrainment, we extended this work using PEA-15 knockout (PEA-15^−/−^) mice to determine how loss of PEA- 15 affects circadian timing, entrainment, and ERK signaling within the SCN. At the cellular level, both clock-gated and light-evoked ERK activation were significantly reduced in PEA- 15^−/−^ mice compared with wild-type controls. Under constant darkness, PEA-15^−/−^ mice exhibited a significantly shorter free-running period relative to wild-type animals.

Assessment of circadian entrainment using jet-lag resetting and Aschoff type I light-pulse paradigms revealed a blunted phase-delay response to light and an accelerated rate of entrainment to a phase-advancing light stimulus. Thus, loss of PEA-15 alters multiple properties of the SCN clock, including its intrinsic period and its responsiveness to photic input, with effects that depend on the direction of phase resetting. Collectively, these findings identify PEA-15 as an important regulator of SCN circadian function and implicate ERK/MAPK signaling as one mechanism through which PEA-15 modulates clock timing and photic entrainment.

## Introduction

Circadian rhythms regulate physiological functions, behaviors, and metabolic activities on an approximate 24-hour schedule. Central to the mammalian circadian timing system is the suprachiasmatic nucleus (SCN) of the hypothalamus, which serves as the principal circadian pacemaker and orchestrates rhythmic processes throughout the body (Mohawk et al., 2012; Moore et al., 2002). The SCN receives photic input from the retina, enabling the circadian system to synchronize with the external light/dark cycle and thereby align internal biological processes with environmental time cues (Ashton et al., 2022; Meijer & Schwartz, 2003). At the molecular level, circadian timing within the SCN is generated by a well-characterized transcriptional-translational feedback loop (TTFL) centered on the core clock proteins BMAL1 and CLOCK. These proteins form a heterodimeric transcription factor that drives the expression of Period and Cryptochrome genes. PER and CRY proteins subsequently accumulate, form complexes, and inhibit BMAL1/CLOCK-dependent transcription, thereby generating the negative-feedback loop underlying circadian oscillations (Takahashi, 2017; Yi et al., 2022). Additional transcriptional regulatory loops and intracellular signaling pathways modulate fundamental properties of the TTFL, including its period, amplitude, and phase, and thereby contribute to the timing properties of the SCN oscillator (Brancaccio et al., 2013; Hastings et al., 2008; O’Neill et al., 2008).

Among these pathways, the p44/42 mitogen-activated protein kinase (ERK/MAPK) pathway has emerged as an important regulator of both circadian timing and photic entrainment (Coogan & Piggins, 2004; Goldsmith & Bell-Pedersen, 2013; Obrietan et al., 1998).

<u>P</u>hosphoprotein <u>E</u>nriched in <u>A</u>strocytes <u>15</u> (PEA-15) is a small, 15-kDa protein originally identified in astrocytes but subsequently shown to be expressed in multiple cell types, including neurons (Wheaton et al., 2018). As a multifunctional adaptor protein lacking intrinsic enzymatic activity, PEA-15 regulates diverse cellular processes, including apoptosis, proliferation, and glucose metabolism (Fiory et al., 2017; Formstecher et al., 2001; Kitsberg et al., 1999; Wei, 2015; Zvalova et al., 2001). A major function of PEA-15 is to serve as a scaffold and anchor for ERK, promoting its sequestration within the cytoplasm and thereby regulating ERK nuclear signaling, and downstream gene transcription (Formstecher et al., 2001; Hill et al., 2002; Vaidyanathan et al., 2007). Through this interaction, PEA-15 has the potential to modulate cellular processes regulated by ERK/MAPK signaling, including those involved in circadian clock function.

Our previous work (Wheaton et al., 2018) demonstrated that PEA-15 is highly expressed within the SCN, with prominent expression in distinct neuronal populations, including arginine vasopressin (AVP)-positive neurons of the SCN “shell” and vasoactive intestinal peptide (VIP)-positive neurons of the “core”. Notably, PEA-15 expression within the SCN exhibits circadian rhythmicity, suggesting that its regulatory functions may themselves be temporally controlled. Photic stimulation during the early subjective night induces phosphorylation of PEA-15 at serine 104 and serine 116, which promotes dissociation of the PEA-15/ERK complex. This light-dependent change in PEA-15/ERK association could facilitate ERK nuclear translocation and downstream signaling events that contribute to photic resetting of the circadian clock (Wheaton et al., 2018).

Although these findings implicate PEA-15 in the regulation of circadian signaling, its functional contribution to SCN clock timing and entrainment remains incompletely understood. In the present study, we investigated the role of PEA-15 in regulating circadian clock function and photic entrainment using PEA-15^−/−^ mice. Using a combination of behavioral assays, protein profiling, and cell-signaling analyses, we demonstrate that loss of PEA-15 alters the intrinsic period of the SCN oscillator and modifies its response to photic stimulation. In parallel, PEA-15 deficiency disrupts both the rhythmic and light-evoked activation of ERK/MAPK signaling within the SCN.

## Methods

### Animals and Experimental Design

#### Transgenic Mouse Model

Mice with the targeted deletion of PEA-15 (PEA-15^−/−^ mice) were initially described in Kitsberg et al. (1999), and are currently available through Jackson Laboratories (Strain #: 030701). The PEA-15^−/−^ line was initially generated in 129S6/SvEvTac-derived TC1 embryonic stem cells, and chimeric males were bred with NIH Black Swiss females. The line was then backcrossed into the C57BL/6J line for a minimum of 10 generations. *PEA*- 15^+/-^ mice were bred to generate PEA-15^−/−^ and WT littermates, which were used for the studies here. Tail-biopsy-based genotyping was performed using either Transnetyx genotyping service (Cordova, TN, USA) or PCR-based profiling using the primer sets and cycling conditions described for Strain # 030701 on the Jackson Laboratories website.

Adult (3-6 months-of-age) male and female mice of each genotype were used in approximately equal numbers in all experiments. No sex-dependent differences were detected in circadian locomotor activity or cellular signaling measures; therefore, data from male and female mice were pooled for subsequent analyses. All animal procedures were approved by the Institutional Animal Care and Use Committee (IACUC) and conducted in compliance with the NIH guidelines.

#### Assessment of Circadian Locomotor Activity

Mice were individually housed in polycarbonate cages equipped with running wheels and maintained under a 12-h light/12-h dark (LD) cycle at an illumination level of 200 lux for at least 14 days before experimental procedures. Circadian locomotor activity was assessed by continuous monitoring of wheel-running behavior. Wheel revolutions were detected using magnetic switches and recorded in 5-min intervals using VitalView software (Starr Life Sciences, Oakmont, PA, USA). ClockLab software (Actimetrics, Lafayette, IN, USA) was used to quantify locomotor activity across the 24-h cycle, including total daily activity, activity during the light and dark phases, and the phase angle of entrainment. For the determination of the phase angle of entrainment, the mean activity-onset time for each animal was calculated relative to the light-to-dark transition (ZT12) over a 10-day recording period. To determine the endogenous free-running period (tau: τ), mice were transferred to constant darkness (DD) for at least 14 days. The free-running period was determined using a χ² periodogram in ClockLab over a 10-day interval beginning after 4 days in DD. ClockLab software was also used to determine average daily activity during DD, expressed as wheel rotations per day, as well as the duration (alpha; α) and acrophase of locomotor activity.

#### Assessment of Light-Induced Phase Resetting

The response of the circadian clock to an acute light stimulus was assessed using an Aschoff Type I phase-shifting protocol (Aschoff, 1965). Mice were transferred from the LD cycle to DD and maintained in constant darkness for 14 days before light stimulation.

Animals were then exposed to white LED light (15-min) at 50 lux at CT15.5 and subsequently returned to DD for at least 10 days. Light-induced phase shifts were determined by linear regression analysis of activity-onset times. A baseline regression line was generated from activity-onset times recorded during the 7 days preceding light exposure, and a second regression line was generated from activity-onset times recorded on days 3–10 following the light pulse. The magnitude of the phase shift was calculated as the difference between the two regression lines on the day of light exposure. Phase shifts are reported as mean ± SEM for each genotype.

#### Assessment of Circadian Re-entrainment

The rate of behavioral re-entrainment was assessed following 8-h advances and delays of the LD cycle. For the advance paradigm, mice were initially entrained to a 2-h light/22-h dark (2:22 LD) schedule. The 2-h light period was then advanced by 8 h, and the timing of activity onset relative to the shifted light period was monitored daily until stable entrainment was achieved. The number of days required to achieve stable entrainment was determined for each animal. Stable entrainment was defined as a minimum of four consecutive days during which activity onset occurred within 15 min following termination of the 2-h light period. A similar protocol was used to assess re-entrainment following an 8- h delay in the LD cycle. Mean daily phase shifts and the time required to achieve stable entrainment were compared between genotypes.

#### Circadian-Time Sampling and Light Stimulation

For experiments examining circadian regulation of ERK activation (pERK), mice were maintained under LD conditions for at least 2 weeks before being transferred to DD for dark adaptation. Animals were maintained in complete darkness for 48 h, with tissue collected on the third day in DD. Circadian pERK expression was assessed at four circadian time points (CT 0; 6; 12; 18) distributed across the 24-h cycle. For experiments conducted under LD conditions, ZT0 was defined as lights-on and ZT12 as lights-off. For experiments requiring tissue collection during the light phase, animals were euthanized between ZT5 and ZT7.

To assess light-evoked ERK activation, mice were exposed to white LED light at 50 lux for 10 min at CT16 (estimated). The estimate of CT16 was based on the preceding LD cycle.

Control animals underwent the same handling procedures but remained in darkness and were not exposed to the light stimulus. Immediately following light exposure, animals were euthanized by cervical dislocation under dim red illumination (<4 lux) generated by an LED array emitting at 650–690 nm. To minimize postmortem photic stimulation during brain removal, the eyes were covered with opaque black tape.

#### Brain Tissue Preparation

Immediately following dissection, brains were placed in chilled, oxygenated physiological saline and sectioned at 500 μm using a VT1200 vibrating microtome (Leica Biosystems, Nussloch, Germany). Tissue sections were fixed for 6 h at room temperature in 4% formaldehyde prepared in PBS (pH 7.4) and subsequently cryoprotected overnight at 4°C in 30% sucrose. Tissue was then sectioned at 40 μm using a freezing microtome for immunohistochemical and immunofluorescence analyses.

#### Immunohistochemical Labeling

For immunohistochemical analysis, tissue sections were incubated in phosphate-buffered saline (PBS; pH 7.4) containing 0.3% hydrogen peroxide (to inhibit endogenous peroxidase activity) and then blocked for 1 h in PBS containing 0.2% Triton X-100 and 0.02% sodium azide and 10% normal goat serum (NGS). Sections were subsequently incubated overnight at 4°C with primary antibodies diluted in PBS containing 0.2% Triton X-100 and 0.02% sodium azide and 5% NGS. The following rabbit primary antibodies were used: PEA-15 (1:9,000; catalog #2780, Cell Signaling Technology, Danvers, MA, USA), phospho-ERK (pERK; 1:3,000; catalog #9101, Cell Signaling Technology) and total ERK (1:1,000; catalog #4695, Cell Signaling Technology).

Following primary antibody incubation, sections were incubated for 2 h at room temperature with a biotinylated anti-rabbit secondary antibody (1:1,000; catalog #BA-1000, Vector Laboratories, San Carlos, CA, USA). Sections were then incubated for 45 min with an avidin-biotin-horseradish peroxidase complex using the Vectastain kit (catalog #PK- 4000, Vector Laboratories). Immunoreactivity was visualized using a nickel-enhanced 3,3′- diaminobenzidine (DAB) substrate (catalog #SK-4100, Vector Laboratories). Sections were rinsed three times between each labeling step, mounted on gelatin-coated slides, and coverslipped using Permount (Fisher Scientific). Images were acquired using a DM IRB microscope (Leica Biosystems).

#### Quantification of Immunoreactivity

Quantitative analysis of immunolabeling was performed using ImageJ software (NIH, Bethesda, MD, USA). For the analysis of circadian clock-gated ERK activation, pERK-positive cells were quantified separately within the core and shell regions of the SCN, with anatomical boundaries defined as illustrated in Fig. 2B. Counts from both SCN nuclei within each section were combined, and measurements from two sections per animal were averaged to generate a single value for each animal.

For analysis of light-evoked ERK activation, pERK-positive cells were quantified across the SCN, with counts from the core and shell regions combined. Two central SCN sections were analyzed for each animal, and values were averaged to generate a single measurement per animal.

ERK immunoreactivity was quantified by densitometric analysis using an inverted 8-bit grayscale ranging from 0 to 255. Mean grayscale values were determined for individual SCN nuclei, averaged within each section, and then averaged across two central SCN sections for each animal. Relative ERK expression was calculated by normalizing values to the mean WT labeling intensity, which was assigned a value of 1. Counting was performed blinded to genotype/condition.

#### Immunofluorescence Labeling and Confocal Imaging

Immunofluorescence labeling was performed to identify neuropeptide-defined regions of the SCN. Tissue sections were incubated in PBS containing 0.2% Triton X-100, 0.02% sodium azide, and 5% bovine serum albumin, followed by overnight incubation at room temperature with antibodies directed against vasoactive intestinal peptide (VIP; 1:250; catalog #RP1108, Boster Biotech) or arginine vasopressin (AVP; 1:200; catalog #403 004, Synaptic Systems). Following primary antibody incubation, sections were treated with Alexa Fluor-conjugated secondary antibodies (1:250; ThermoFisher, Waltham, MA USA) specific to the primary antibodies used. Nuclear labeling was performed by incubating sections with Hoechst 33342 (1:1,000; catalog #62249, ThermoFisher) for 20 min. Tissue was washed four times in PBS between labeling steps, mounted on glass slides, and coverslipped using Fluoromount-G (catalog #0100-01, SouthernBiotech, Birmingham, AL, USA). Fluorescence images were acquired using a Leica SP8 confocal microscope. Image-acquisition parameters were maintained consistently across samples and experimental conditions within each fluorescence channel, including pinhole size and contrast settings.

#### Statistical Analysis

Statistical analyses were performed using GraphPad Prism software. The animal was considered the biological replicate for all statistical analyses; measurements obtained from multiple sections or SCN nuclei from the same animal were averaged before statistical testing. Behavioral measures were compared between genotypes using Student’s *t*-test, unless otherwise indicated. Comparisons of clock-gated and light-evoked pERK expression were performed using two-way analysis of variance (ANOVA), and, when appropriate, the analysis was followed by Bonferroni post hoc multiple-comparison tests.

ERK and MEK expression levels and Aschoff type I light pulse data were compared between genotypes using Student’s *t*-test. Analysis of the rate of re-entrainment in the 2:22 light/dark paradigm, which involves non-normally distributed data, were compared using the Mann–Whitney U test. Data are presented as mean ± SEM, and statistical significance was defined as *p* < 0.05.

## Results

### PEA-15 deletion and ERK signaling in the SCN

To investigate PEA-15 signaling in the SCN, we used a genetically modified mouse model lacking the PEA-15 gene (Fig. 1A). To verify that gene deletion resulted in the absence of PEA-15 protein expression, we performed immunohistochemical labeling for PEA-15 on coronal brain sections containing the SCN from wild-type (WT) and PEA-15 knockout (PEA- 15^−/−^) mice. As illustrated in Figure 1B, PEA-15 expression was detected in the SCN of WT mice, with particularly strong expression in the dorsal and lateral regions of the SCN. In contrast, PEA-15 protein was not detected in brain sections from PEA-15^−/−^ mice.

**Figure 1.**
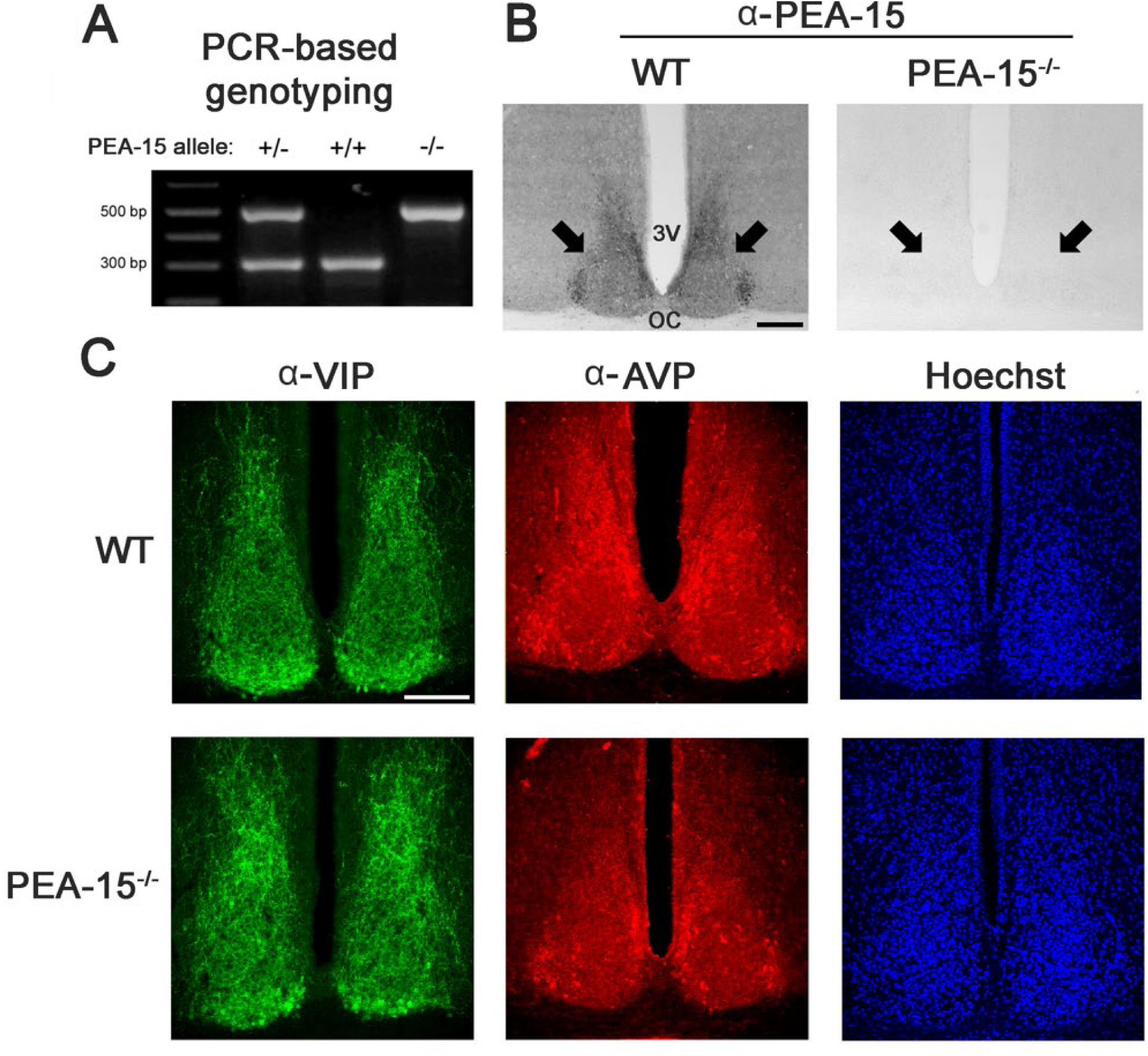
Targeted deletion of *PEA-15* eliminates PEA-15 protein expression without altering gross SCN organization. (A) Representative PCR-based genotyping of mice carrying the wild-type (+/+), heterozygous (+/−), or targeted (−/−) *PEA- 15* allele. (B) Representative coronal sections through the SCN from WT and *Pea15*^-/-^ mice that were labeled (via DAB-based immunohistochemistry) for PEA-15 (α-PEA-15). Robust PEA-15 immunoreactivity was detected throughout the SCN of WT mice, with particularly prominent labeling in the dorsal and lateral regions, whereas PEA-15 immunoreactivity was absent in *PEA-15*^-/-^ mice. Arrows denote the location of the SCN. 3V: Third Ventricle; OC: Optic Chiasm. (C) Representative immunofluorescence images showing the distribution of vasoactive intestinal peptide (VIP; green) and arginine vasopressin (AVP; red) in the SCN of WT and *PEA-15*^-/-^ mice. VIP immunoreactivity was concentrated in the ventral SCN and extended dorsally through extensive dendritic arbors, whereas AVP immunoreactivity was predominantly localized to the dorsal and lateral SCN. The overall distribution of VIP- and AVP-positive neurons was comparable between genotypes. Nuclei are counterstained with Hoechst (blue). Scale bar in *B* and *C*: 250 microns.

**Figure 2.**
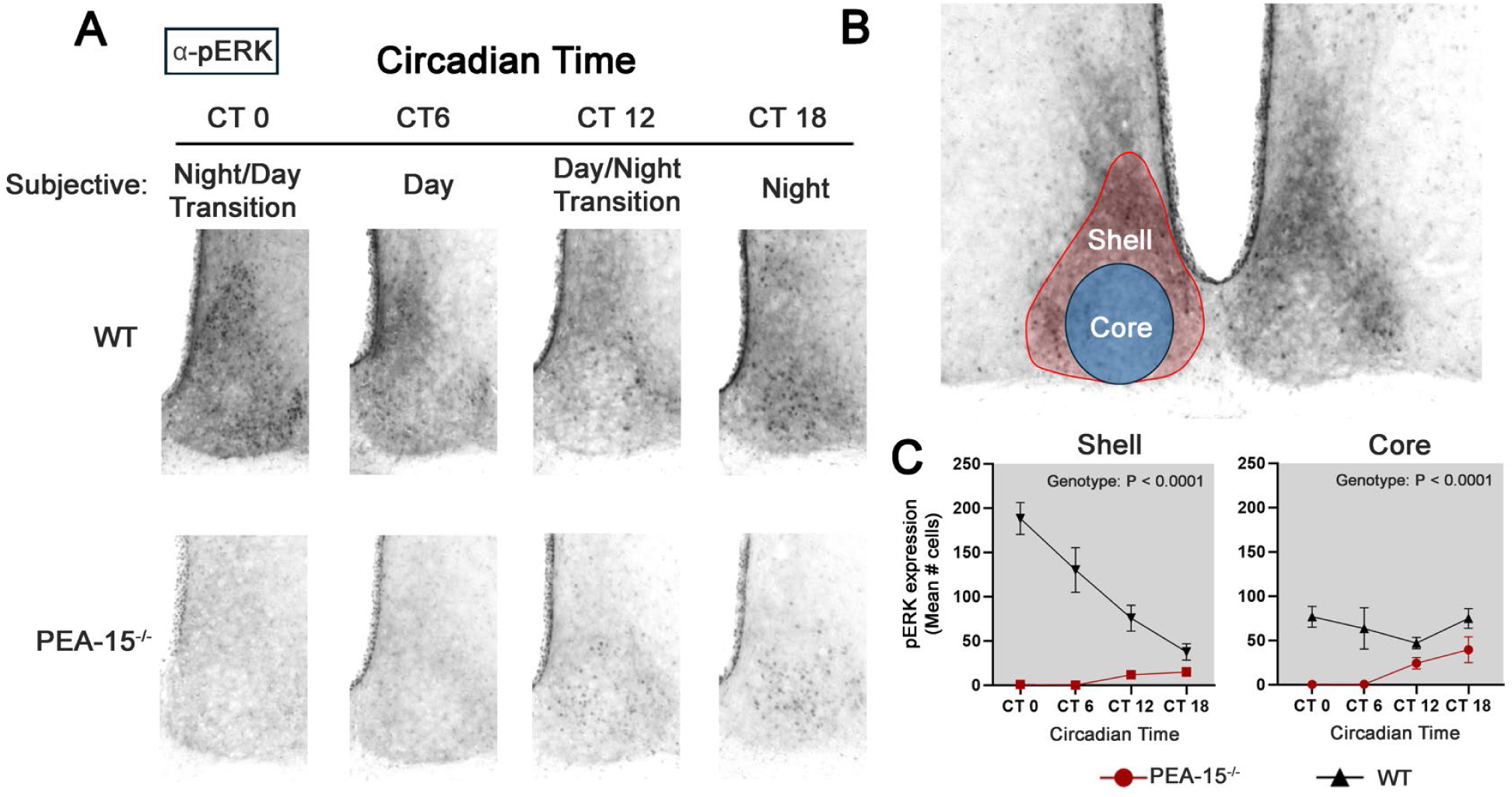
PEA-15 is required for circadian regulation of ERK activation in the SCN. (A) Representative immunohistochemical images showing expression of activated ERK (α-pERK; dually phosphorylated ERK1/2 in the SCN) of WT and PEA-15^-/-^ mice sampled at four circadian times (CT0, CT6, CT12, and CT18) following 2 days of dark adaptation. In WT mice, pERK expression exhibited the expected circadian pattern, with robust activation throughout the SCN shell during the subjective day and comparatively limited activation within the SCN core. In contrast, PEA-15^-/-^ mice exhibited a marked disruption of the circadian pERK expression pattern within the shell, and reduced core pERK across the sampled circadian times. (B) Representative SCN image illustrating the anatomical boundaries used to define the shell and core regions for quantitative analysis. (C) Quantification of pERK expression across the circadian cycle within the SCN shell (left) and core (right) of WT and PEA-15^-/-^ mice. Data are shown as mean ± SEM, and analyzed via two-way ANOVA. The noted “P values” indicate the main effect of genotype, averaged across circadian times; they do not denote comparisons at individual time points.

Immunolabeling-based profiling indicated that the overall organization of the SCN appeared grossly normal in PEA-15^−/−^ mice. Consistent with this observation, the distribution of two well-characterized peptidergic neuronal markers of the SCN, vasoactive intestinal peptide (VIP) and arginine vasopressin (AVP), was similar in PEA-15^−/−^ and WT mice (Fig. 1C). In both genotypes, VIP-expressing neurons were concentrated in the ventral region of the SCN, referred to as the SCN ‘core’, with VIP-positive dendritic arbors extending dorsally from the cell bodies throughout the SCN. Similarly, AVP expression was predominantly observed in neurons within the dorsal and lateral regions of the SCN, referred to as the SCN ‘shell’, in both PEA-15^−/−^ and WT mice (please refer to figure 2B for a depiction of the SCN ‘core’ and ‘shell’).

At the molecular level, PEA-15 functions as a scaffold for ERK, the primary effector of the p44/42 mitogen-activated protein kinase (MAPK) pathway. By binding directly to ERK, PEA- 15 regulates the duration of ERK activation and its capacity for nuclear translocation (Formstecher et al., 2001; Mace et al., 2013). Notably, previous work has shown that ERK activation is modulated across the circadian cycle and that photic input drives robust ERK activation during the nighttime (Obrietan et al., 1998). Given these findings, we examined clock- and light-evoked ERK activation in PEA-15^−/−^ mice.

To characterize ERK activation across the circadian cycle, mice were initially dark-adapted for 2 days, after which SCN tissue was collected at four circadian time (CT) points (CT 0, 6, 12, and 18) and processed immunohistochemically for the activated, dually phosphorylated form of ERK (pERK). Previous work has revealed a complex spatial and temporal pattern of ERK activation in the SCN across the circadian cycle. Along these lines, Obrietan et al. (1998) reported that pERK expression in the shell region is observed primarily during the subjective day. In contrast, only limited subsets of neurons within the core express activated ERK across the circadian cycle.

Consistent with this, here we show that in WT mice, pERK immunoreactivity is prominent within the shell during the subjective day, with comparatively lower levels in the core (Fig. 2A). To assess the effects of PEA-15 loss on this temporal pattern, pERK-immunopositive cells were quantified separately in the core and shell at CT 0, 6, 12, and 18, using each animal’s average count per section (n = 3 independent animals per genotype per time point). Separate two-way ANOVAs revealed significant main effects of genotype in both the core (F(1,16) = 35.42, P < 0.0001) and shell (F(1,16) = 127.25, P < 0.0001), with lower counts in PEA-15^−/−^ mice when averaged across the sampled times. Within the shell, WT mean counts decreased across the circadian cycle (from the CT 0 to CT 18), whereas counts in PEA-15^−/−^ mice were near zero at CT 0–6 and higher at CT 12–18 (Fig. 2A–C). Both the main effect of circadian time (F(3,16) = 10.75, P = 0.0004) and the genotype × time interaction (F(3,16) = 16.38, P < 0.0001) were significant, supporting an altered temporal profile of shell pERK expression following PEA-15 loss. In the core, pERK-immunopositive cells remained readily detectable in PEA-15^−/−^ mice at CT 12 and CT 18 despite the overall reduction in counts (Fig. 2A–C). However, neither the main effect of circadian time (F(3,16) = 1.83, P = 0.182) nor the genotype × time interaction (F(3,16) = 2.20, P = 0.128) was significant, providing insufficient evidence for a genotype-dependent difference in the core temporal profile.

To examine light-evoked ERK activation, we utilized a well-characterized light-pulse paradigm in which mice were initially dark-adapted for 2 days and then exposed to light (50 lux) for 10 min during the circadian night (CT 16). Mice were immediately sacrificed following the light pulse, and SCN tissue was processed for pERK expression. In WT mice, this light-treatment paradigm resulted in robust ERK activation. Interestingly, however, the magnitude of light-evoked ERK activation was significantly reduced in PEA-15^−/−^ mice relative to WT mice (Fig. 3A & 3B).

**Figure 3.**
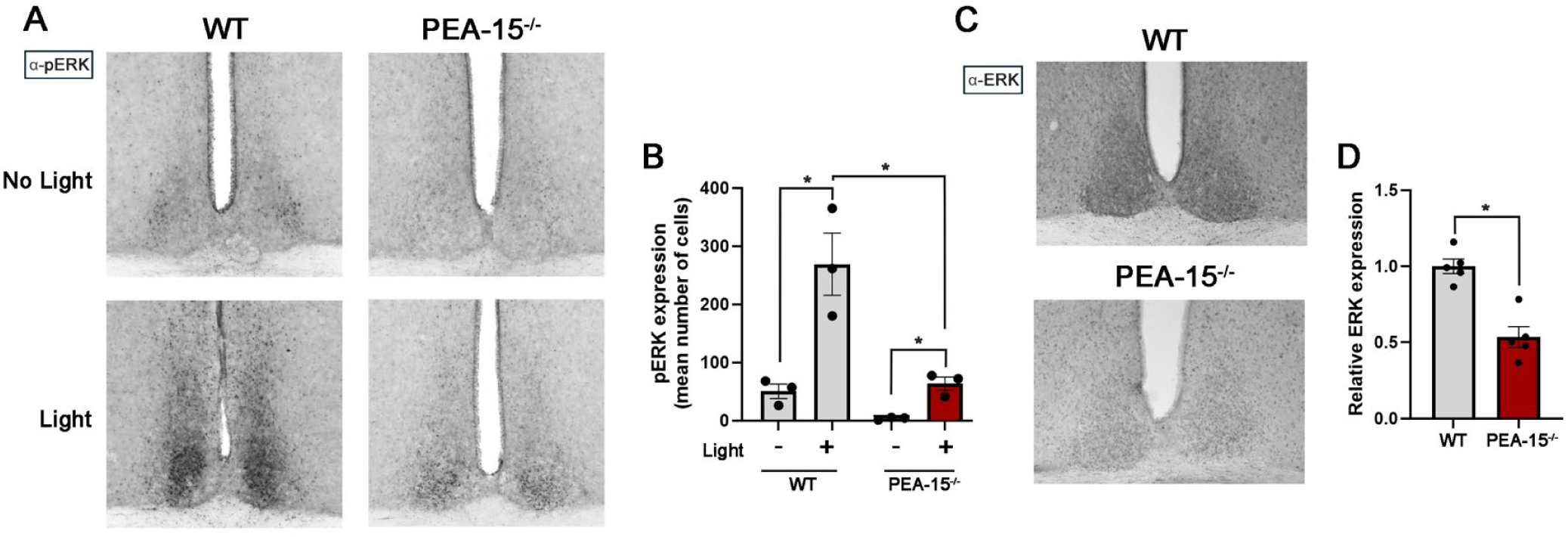
Light-evoked ERK activation and ERK expression in the SCN. (A) Representative images of pERK immunoreactivity in the SCN of wild-type (WT) and PEA-15^-/-^ mice under dark-adapted baseline conditions (No light) or following exposure to a 10-min light pulse (50 lux) at CT16. Light exposure produced a robust increase in pERK expression throughout the SCN of WT mice, whereas light-evoked ERK activation was markedly attenuated in PEA-15^-/-^ mice. (B) Quantification of pERK expression in the SCN under baseline and light-pulse conditions. Light significantly increased pERK expression in both genotypes, with the magnitude of light-evoked ERK activation significantly reduced in PEA- 15^-/-^ mice compared with WT mice. (C) Representative images of total ERK immunoreactivity in the SCN of WT and PEA-15^-/-^ mice collected during the middle of the light phase (ZT5–7). (D) Quantification of relative total ERK expression in the SCN demonstrating significantly reduced ERK protein levels in PEA-15^-/-^ mice compared with WT mice. Data are presented as mean ± SEM, with individual data points shown. Statistical significance for panel *B* was determined by two-way ANOVA followed by a Bonferroni multiple-comparisons test; *p* < 0.01. Statistical significance for panel *D* was determined using an unpaired Student’s *t*-test; * *p* < 0.01.

Although PEA-15 is best characterized as a modulator of ERK phosphorylation kinetics and subcellular localization, previous studies suggest that PEA-15 may also influence ERK protein stability and expression (Formstecher et al., 2001; Mace et al., 2013). To investigate the potential effects of PEA-15 deletion on total ERK (ERK1 and ERK2) protein levels, we performed immunohistochemical analysis of ERK expression in SCN tissue collected during the middle of the day, zeitgeber time (ZT) 5–7 (Fig. 3C). ERK expression was significantly decreased in PEA-15^−/−^ mice compared with WT mice (Fig. 3D). This reduction in expression was observed throughout the dorsoventral axis of the SCN. These findings indicate that PEA-15 is required to maintain normal physiological levels of ERK within the SCN, suggesting that loss of this scaffold protein may predispose ERK to accelerated turnover.

### PEA-15 deletion alters circadian timing and photic entrainment

Next, we examined the effects of PEA-15 deletion on the circadian timing properties of the SCN. To this end, we used a wheel-running activity paradigm as a functional readout of the intrinsic timing and entrainment properties of the circadian system in PEA-15^−/−^ mice and WT littermates. Animals were initially entrained to a standard 12-h light/dark (LD) cycle for 10 days, followed by transfer to constant darkness (DD).

Both genotypes successfully entrained to the LD cycle and exhibited the expected nighttime bout of locomotor activity (Fig. 4A). Interestingly, however, total locomotor activity was significantly lower in PEA-15^−/−^ mice than in WT animals (Fig. 4B). Analysis of running activity as a function of time of day showed that PEA-15^−/−^ mice exhibited significantly less activity during the nighttime compared with WT mice. However, the day/night activity ratio did not differ significantly between genotypes (Fig. 4C-E). With respect to entrainment to the 12-h LD cycle, PEA-15^−/−^ mice did not exhibit a difference in the phase angle of entrainment (based on the onset of locomotor activity) compared with WT animals (Fig. 4F).

**Figure 4.**
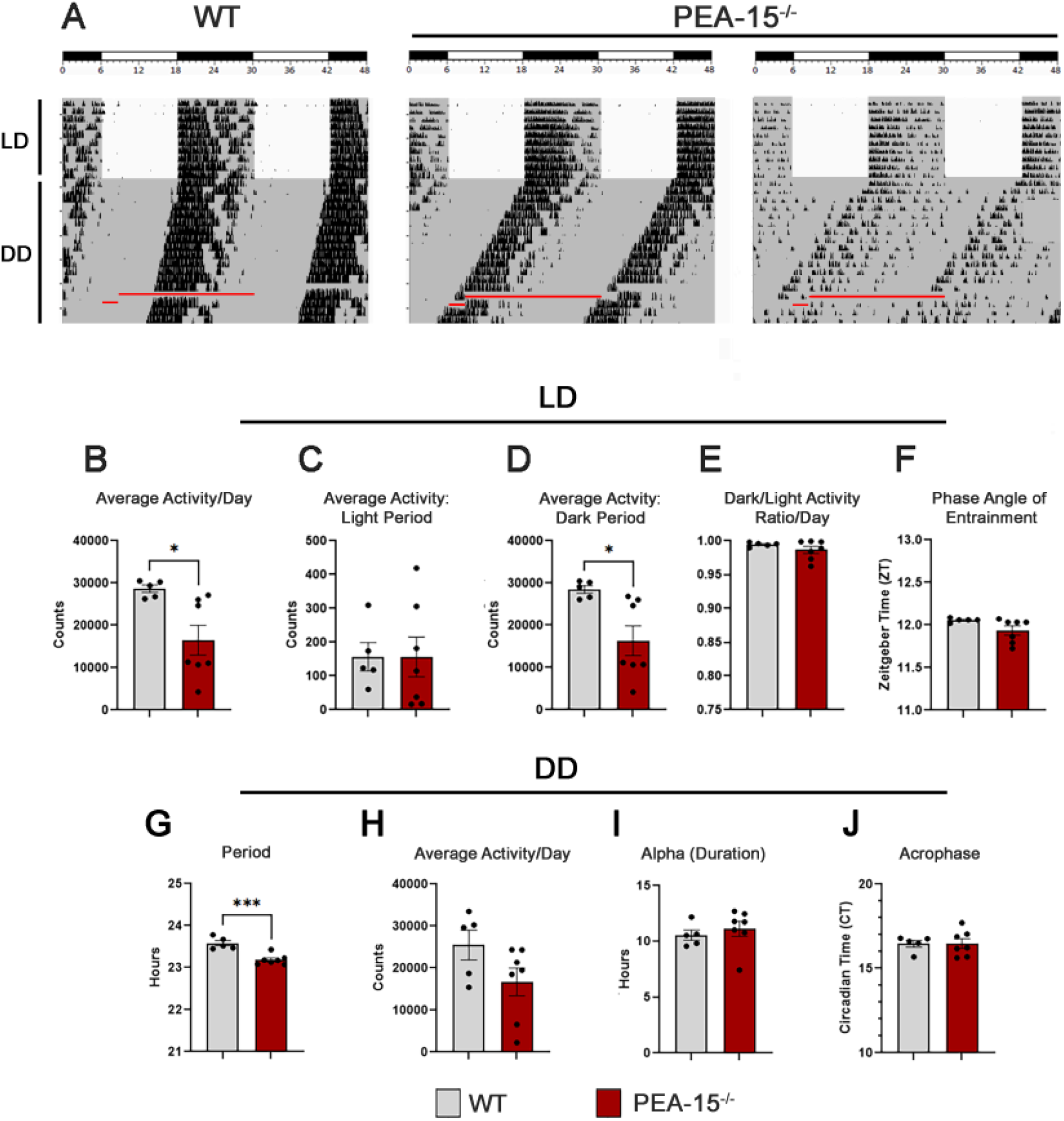
PEA-15 deletion reduces locomotor activity and shortens the intrinsic circadian period. (A) Representative double-plotted actograms from one WT and two PEA- 15^−/−^ mice under LD and DD conditions. Black vertical marks indicate wheel-running activity; white and gray backgrounds indicate light and dark intervals, respectively. The single-plotted red horizontal lines indicate periods during which behavioral data were not collected. (B–F) Quantification of average daily activity (B), activity during the light phase (C), activity during the dark phase (D), the proportion of total daily activity occurring during the dark phase (E), and the phase angle of entrainment, expressed as the Zeitgeber Time (ZT) of activity onset (F). PEA-15^−/−^ mice exhibited significantly lower total daily and dark-phase activity, whereas light-phase activity, the proportion of dark-phase activity, and activity onset did not differ significantly between genotypes. (G-J) Quantification of average free-running circadian period (τ; G), daily activity (H), duration of the daily active interval (alpha; I), and acrophase, expressed in circadian time (CT; J). PEA-15^−/−^ mice exhibited a significantly shorter free-running period, with no significant differences in daily activity, alpha, or acrophase. Individual points represent individual animals. Statistical significance was determined using an unpaired Student’s *t*-test; *P < 0.05; ***P < 0.001.

To examine intrinsic circadian timing properties, PEA-15^−/−^ mice were placed in constant darkness (Fig. 4A). Under these conditions, the intrinsic period (τ) of the circadian activity rhythm was significantly shorter in PEA-15^−/−^ mice than in WT counterparts (Fig. 4G), indicating that PEA-15 contributes to the regulation of circadian period. No significant differences in the average daily activity, the duration of daily activity (alpha) or acrophase were observed between PEA-15^−/−^ and WT mice (Fig. 4H-J).

To examine the contribution of PEA-15 to the light-entrainment properties of the circadian system, we utilized a series of light-evoked phase-shifting paradigms. In the first paradigm, mice were initially entrained to a 2:22 light/dark cycle (2 h light/22 h dark), after which the light cycle was either advanced or delayed by 8 h, and the rate of re-entrainment was determined (Fig. 5A). Initially, both WT and PEA-15^−/−^ mice stably entrained to the 2:22 light/dark cycle, with a marked bout of locomotor activity occurring following the transition from light to dark. Interestingly, PEA-15^−/−^ mice required significantly more time to stably entrain following an 8-h delay in the light cycle compared with WT mice. The majority of WT mice (5/6 mice) required approximately 3 days to re-entrain to the new light cycle. In contrast, the majority of PEA-15^−/−^ mice (5/6 mice) required over 3X longer (≥ 9 days) to achieve stable entrainment to the delayed light cycle (Fig. 5B). Interestingly, two PEA-15^−/−^ mice responded to the delaying phase shift by phase-advancing until their activity periods aligned with the 2-h light period (an example is provided in Fig. 5A: far right side actograph). In contrast to the prolonged re-entrainment observed following an 8-h delay, PEA-15^−/−^ mice re-entrained significantly faster following an 8-h advance in the light cycle (Fig. 5A & 5C), achieving stable entrainment approximately 40% faster than WT mice (15 vs. 24 days, respectively).

**Figure 5.**
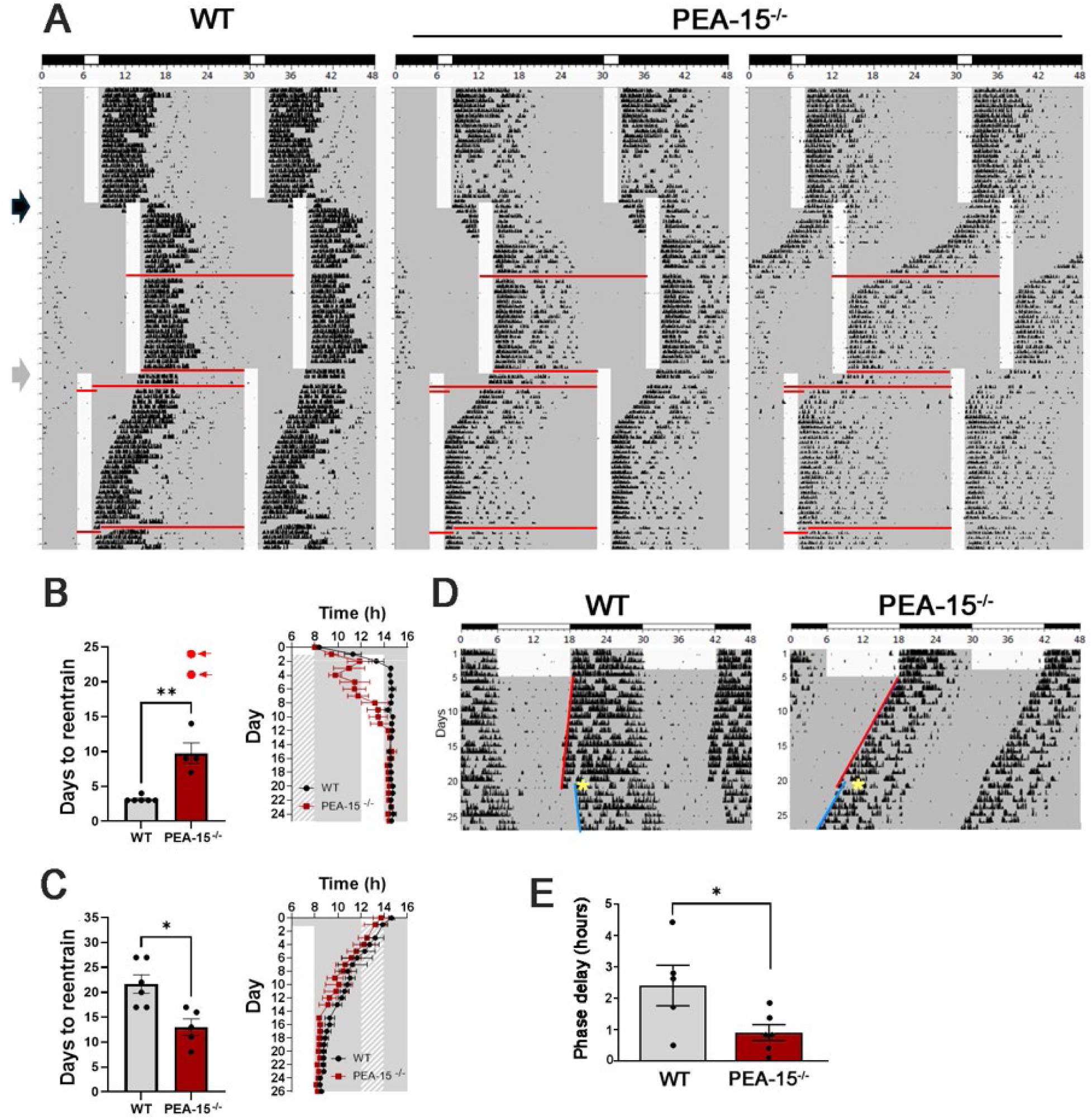
PEA-15 regulates photic resetting and re-entrainment of the circadian clock. (A) Representative double-plotted actograms from WT and PEA-15^−/−^ mice maintained on a 2-h light/22-h dark (2:22 LD) cycle and subjected to 8-h delays and advances of the light cycle. White vertical bars indicate the 2-h light period, and single-plotted red horizontal lines indicate intervals during which behavioral data were not collected. Arrows indicate the onset of the 8-h phase delay (black) and phase advance (gray). The PEA-15^−/−^ mouse shown at right responded to the 8-h delay by phase-advancing its activity rhythm until it aligned with the shifted light cycle. (B, C) Number of days required to achieve stable re- entrainment following an 8-h delay (B) or advance (C) of the light cycle. PEA-15^−/−^ mice re- entrained significantly more slowly following the phase delay but significantly more rapidly following the phase advance compared with WT mice. Two PEA-15^−/−^ mice responded to the phase-delay paradigm by phase-advancing rather than delaying; their re-entrainment times are indicated by red arrows in B and were not included in the statistical comparison. Data were analyzed using the Mann–Whitney U test. Corresponding plots at right show the mean daily phase resetting of WT and PEA-15^−/−^ mice following the 8-h shifts. White rectangles denote the timing of the light period before the shift (Day 1) and after the shift (Days 2–26); cross-hatched rectangles indicate the timing of the light period prior to the shift. (D) Representative double-plotted actograms from dark-adapted WT and PEA-15^−/−^ mice exposed to a 15-min, 50-lux light pulse at CT15.5. Red and blue regression lines indicate the phase of the activity rhythm before and after the light pulse, respectively. (E) Quantification of light-induced phase delays. PEA-15^−/−^ mice exhibited significantly smaller phase delays than WT mice. Data were analyzed using an unpaired Student’s *t*-test. Data are presented as mean ± SEM, with individual points representing individual animals. *P* < 0.05; P < 0.01.

To further probe the responsiveness of the circadian system to photic input, we used an Aschoff Type I resetting paradigm in which a brief light pulse (15 min, 50 lux) was delivered at CT 15.5 (early subjective night). This phase-delaying stimulus elicited a significantly smaller phase shift in PEA-15^−/−^ mice than in WT animals, representing a greater than two-fold reduction in the magnitude of the phase shift (Fig. 5D & 5E). These findings indicate that PEA-15 modulates the photic resetting response of the circadian timing system.

## DISCUSSION

While the central components of the TTFL that generates circadian rhythms are well characterized, the signaling pathways that modulate these loops and integrate environmental entrainment cues remain less well understood. Here, we demonstrate that PEA-15 is a critical regulator of SCN clock timing and photic entrainment. Genetic deletion of PEA-15 resulted in a shorter free-running period (τ) under constant darkness, a severely blunted phase-delaying response to early-night light pulses and accelerated re-entrainment to a phase-advancing stimulus. Biochemically, these phenotypes were accompanied by a loss of daytime clock-gated ERK phosphorylation rhythms in the SCN shell, reduced light-evoked ERK activation, and an approximately 50% reduction in total ERK protein. Together, these findings identify PEA-15 as an important regulator of the SCN timing system and suggest that its effects on circadian period and photic entrainment are mediated, at least in part, through modulation of ERK/MAPK signaling.

### PEA-15 and ERK Expression and Activation in the SCN

Notably, with respect to ERK stability, the absence of PEA-15 likely predisposed ERK to accelerated turnover and degradation, which would be expected to contribute to the reduction in total SCN ERK expression. Further, with respect to ERK activity, in PEA-15^−/−^ mice there is nearly a complete loss of endogenous clock-gated ERK activity rhythms during the daytime. Although the noted reduction in ERK expression would contribute to this reduction in rhythmic activity, reduced expression cannot fully account for the loss of its daytime activation. Instead, this severe signaling deficit likely reflects additional molecular alterations. First, deleting PEA-15 results in a loss of critical scaffolding that facilitates kinase coupling between upstream MEK and ERK (Ramos et al., 2000). In a functional MAPK signaling module, PEA-15 acts to coordinate these kinases, ensuring high-affinity interactions and rapid signal transduction; Without this spatial organization, MEK- to-ERK kinase coupling efficiency would likely be markedly reduced. Second, the loss of PEA-15 impairs the stability of the phosphorylated form of ERK. PEA-15 is known to physically bind dually phosphorylated ERK, creating a steric shield that protects the activated kinase from dephosphorylation by dual-specificity phosphatases (DUSPs) and MAP kinase phosphatases (MKPs) (Mace et al., 2013). This shielding effect normally lengthens the activation period of phosphorylated ERK. In the absence of PEA-15, activated ERK is likely to be more effectively targeted for dephosphorylation, and this, in turn, could result in a dramatic shortening of the ERK active state duration. Clearly our data cannot provide a specific mechanism by which PEA15 deletion leads to the profound effects on clock-gated ERK activity reported here. Nevertheless, the noted cellular and molecular based studies provide potential mechanisms that contribute to the marked disruption in ERK activity rhythms.

The reduction in ERK signaling was not limited to endogenous clock-gated activity. PEA- 15^−/−^ mice also exhibited a reduction in light-evoked ERK activation, a finding of particular significance given the established role of the MAPK/ERK pathway in coupling photic input to circadian entrainment. Light exposure during the subjective night induces rapid ERK phosphorylation in the SCN and activates downstream transcriptional pathways, including CREB and Elk-1 (Coogan & Piggins, 2003; Obrietan et al., 1998). These signaling events promote the expression of immediate-early genes and clock genes such as Period1 (Butcher et al., 2005) thereby translating photic input into phase-dependent resetting of the molecular oscillator (Butcher et al., 2002; Coogan & Piggins, 2003). Within the SCN, ERK/MAPK signaling also functions as an upstream activator of mTOR-mediated translation, providing an additional mechanism through which photic stimulation can influence clock-gene expression (Cao et al., 2008; Cao et al., 2010). Consistent with this established role of ERK/MAPK signaling, the reduction in light-evoked ERK activation in PEA-15^−/−^ mice was accompanied by markedly attenuated phase delays in both the jet-lag phase-delay paradigm and the Aschoff type I light-pulse paradigm.

Interestingly however, the rate of reentrainment to an 8 hr advance in the light cycle was accelerated in the PEA-15^−/−^ mouse line. There are several potential reasons for this accelerated rate of reentrainment. One possibility is that the shorter τ in the PEA-15^−/−^ mice (discussed below) would allow for more rapid reentrainment to a phase-advancing stimulus. Clearly additional experiments will be needed to address these potential mechanisms.

### Disruption of Clock-Gated ERK Rhythms and SCN Period

In wild-type mice, SCN neurons display coordinated, clock-gated rhythms of pERK activation that are particularly prominent in the AVP-positive shell (Obrietan et al., 1998). In PEA-15^−/−^ mice, this endogenous rhythm was severely disrupted, with shell pERK expression failing to reach normal peak levels. Interestingly, the near-complete loss of daytime pERK rhythmicity coincided with a significant shortening of the period (τ) of the circadian oscillator.

While direct evidence connecting the loss of pERK rhythms to a shortened τ is lacking, there are, nevertheless several established cellular- and circuit-based mechanisms that could explain how the disruption of clock-gated ERK signaling could contribute to the shortened τ observed in PEA-15^−/−^ mice. These include changes in 1) the coupling architecture of the SCN network, 2) altered translational or post-transcriptional regulation of core clock and clock-controlled proteins, and 3) disruption of the daily rhythm of neuronal excitability. Importantly, these mechanisms are not necessarily independent, because ERK signaling is positioned to influence multiple intracellular and intercellular processes simultaneously.

With respect to the idea that PEA-15/ERK signaling could affect clock neuron network coupling (point 1), the period of the SCN results from interactions among thousands of cellular clocks, and the ERK/MAPK pathway acts downstream of paracrine systems that set this period, including VIP and AVP signaling. The loss of rhythmic ERK activity in PEA-15^−/−^ mice could, therefore, disrupt how these signals coordinate the timing of discrete cellular oscillator populations. Such a disruption could alter the ensemble period by reducing phase coherence among neurons with different intrinsic periods or by shifting the relative contributions of faster and slower oscillator populations. This idea is supported by data showing that the overexpression of the ERK phosphatase DUSP4 in SCN neurons shortens the circadian period, while targeted DUSP4 expression in Vpac2-expressing neurons attenuates the period-lengthening effects of VIP (Hamnett et al., 2019). Of note, microdissection studies have shown that the intrinsic period of the AVPergic dorsal SCN is shorter than the period of the ventral SCN (Noguchi et al., 2004), thus indicating that core/shell communication is central to setting the overall periodicity of the SCN. Thus, the loss of rhythmic MAPK activity in AVP-expressing neurons raises the possibility that MAPK signaling in the shell could underlie core/shell signaling, and in turn, the intrinsic timing properties of the SCN.

With respect to the idea that PEA-15/ERK signaling could affect the translational and post-transcriptional regulation of core clock and clock-gated genes (point 2), support for this idea comes from work showing that ERK has a direct effect on core clock machinery. Along these lines, ERK mediated phosphorylation of BMAL1 has been shown to negatively regulate CLOCK:BMAL1-mediated transcriptional activation, providing a potential regulatory brake within the TTFL (Sanada et al., 2002). In PEA-15^−/−^ mice, the loss of clock-gated pERK would likely relieve this BMAL1-directed inhibition, and, absent this brake, the CLOCK:BMAL1 complex could exhibit accelerated transactivation kinetics, leading to faster accumulation of *Per* and *Cry* transcripts and a subsequent shortening of the free-running period of the oscillator.

Finally, with respect to point 3, the molecular clock and neuronal excitability are bidirectionally coupled in the SCN: membrane activity and associated calcium signaling help sustain and synchronize clock-gene rhythms, while molecular clock-dependent mechanisms regulate the ionic conductance and cellular excitability underlying rhythmic electrical activity (Harvey et al., 2020; Lundkvist et al., 2005; Yamaguchi et al., 2003).

Within the context of our data, it is interesting to note that the pharmacological inhibition of MAPK signaling damps circadian clock-gene expression and attenuates both evoked and autonomous circadian neuronal activity (Akashi et al., 2008; Dziema & Obrietan, 2002).

Further, ERK signaling regulates A-type potassium currents (Adams et al., 2000; Schrader et al., 2006), which are important determinants of neuronal excitability, and have been shown to affect SCN clock timing and entrainment (Granados-Fuentes et al., 2012).

Clearly, the three potential mechanisms outlined here are speculative-however, they provide a testable framework for understanding how MAPK signaling could play a central role in setting the period of the SCN oscillator.

